# Nectar Sugar Enhancement in Response to Bee Buzzing in *Rhododendron* × *pulchrum*: Sound-sensing Organs and Sensitivity Range

**DOI:** 10.1101/2025.09.09.675125

**Authors:** Kokomi Seike, Atsushi Tani, Gaku S. Hirayama, Atushi Ushimaru

## Abstract

Plants can perceive and respond to airborne sounds, but our current understandings of this phenomenon remain limited. Flowers are known to exhibit increased nectar sugar concentration in response to pollinator sounds; however, there has been only a single report from *Oenothera drummondii*. To test the generality of pollinator-sound-induced nectar enrichment in bee-pollinated plants, we examined nectar responses to airborne sounds, particularly those produced by bee buzzing, in *Rhododendron* × *pulchrum* and *Lamium amplexicaule*. We exposed the flowers to playback sounds of bee buzzing, a similar synthetic sounds signal (200 Hz) and those with higher frequency (5000 Hz) and compared nectar sugar concentrations among these sound-treated and silent control flowers. We found an approximately 10% increase in nectar sugar concentration in response to playback sounds of bee buzzing and synthetic sounds signals at bee-like frequency (200 Hz), while no such response was observed under higher-frequency stimuli or silent condition in *R*. × *pulchrum*. A similar response to bee sounds was also observed in *L. amplexicaule*. Furthermore, these responses were observed at a sound pressure level of 65 or 100 dB, but not at 50 dB in *R*. × *pulchrum*, indicating a sound sensitivity threshold. We experimentally removed petals, stamens, or pistils (alone or in combination) from flowers and exposed them to 200 Hz sounds and silent treatments and found that both petals and stamens were necessary for such sound responses, suggesting that these floral organs are involved in acoustic sensing. Our findings provide new insights into floral sound sensing and suggest that acoustic responses to pollinator visitations may be more widespread among bee-pollinated flowering plants.

## Introduction

Although plants lack specialized auditory organs or nervous systems, they can perceive and respond to sound stimuli (Jung *et al*. 2018; Appel *et al*. 2014). Recent studies have demonstrated that exposure to specific acoustic frequencies can influence plant physiology, whereby specific sound stimuli promote growth or alter gene expression and hormone levels in some plant species (Ghosh *et al*. 2016; Ghosh *et al*. 2017; Hassanien *et al*. 2014).

Flowers can also actively respond to sounds generated by their specific pollinators. Among such sound-related flower responses, pollen release induced by pollinator-generated vibrations has been a traditionally known phenomenon in various flowering plant taxa (Buchmann, 1978; Pritchard & Vallejo-Marín, 2020). In buzz-pollinated species such as *Solanum* species, specialized bee-produced vibrations within the 100–400 Hz range mechanically stimulate anthers to explosively release pollen, thereby avoiding pollen thieves by ineffective pollinators (King & Buchmann, 1996; De Luca & Vallejo-Marin, 2013; Rosi-Denadai *et al*. 2018).

Moreover, Veits *et al*. (2019) recently reported that flowers of bee- and hawkmoth-pollinated *Oenothera drummondii* increased their nectar sugar concentration within three minutes when exposed to bee buzzing sounds. The induced nectar concentration increase would be adaptive because bees have been shown to be capable of perceiving differences in sugar concentrations, as small as 1-3% (Mujagic *et al*. 2009; Whitney *et al*. 2008), and to selectively prefer flowers with higher sugar concentrations. These findings suggest that some plants are capable of perceiving pollinator-related sounds and adjusting their floral rewards to enhance pollination success.

However, nectar sugar enhancement in response to pollinators’ sounds has been demonstrated in only a single species, and therefore, the generality of this phenomenon in angiosperms remains to be tested. Moreover, the auditory threshold of sound intensities to which flowers are responsive remains unknown, and the specific floral organs responsible for acoustic perception have not been sufficiently examined, although Veits *et al*. (2019) suggested that the petals may serve as sound-receiving structures based on vibrational responses.

In this study, we investigated the effects of short-term buzzing sounds exposure on nectar sugar concentration in bee-pollinated (melittophilous) *Rhododendron* × *pulchrum* and *L. amplexicaule* under field conditions. We examined the following three questions. First, is nectar sugar concentration enhanced by bee buzzing and/or artificial sounds in both *R*. × *pulchrum* and *L. amplexicaule*? Second, does an acoustic threshold for sound response exist in *Rhododendron* flowers? Third, what floral organs are responsible for sound sensing in *Rhododendron* flowers? By answering these questions, we discuss the generality and adaptive significance of buzzing-induced nectar quality enhancement in melittophilous flowers.

## Material and Methods

### Study species and sites

We examined two bee-pollinated species, *R*. × *pulchrum* and *L. amplexicaule* at Aotani-cho park and Kobe University in Hyogo Prefecture, Japan. *Rhododendron* × *pulchrum* (Hirado azalea, Ericaceae) is an ornamental shrub species, which is a natural hybrid of several *Rhododendron* species that have been cultivated in Hirado, Nagasaki Prefecture, Japan (Shen *et al*. 2022). The species is commonly planted globally in parks and blooms from April to May. *Rhododendron* × *pulchrum* produces large, funnel-shaped, long-tubed, and pink flowers, which are pollinated mainly by several kinds of bees including bumble bees, honey bees, and Anthophorinae and Eucerini species in the study sites.

*Lamium amplexicaule* (Lamiaceae) is an annual herb that is natively distributed in temperate Asia, Europe, and north Africa, often growing around rice fields and along roadsides. From March to June, this species produces pink long-tubed chasmogamous flowers which are visited by Anthophorinae and Eucerini species as well as cleistogamous flowers.

### Field experiments

#### Sound treatments

We experimentally prepared 2–4 types of flowers exposed to different acoustic stimuli: (1) those exposed to buzzing sounds recorded from a honey beehive (bee treatment); (2) those exposed to artificial sinusoidal tones at 200 Hz (200 Hz treatment); (3) those exposed to artificial sinusoidal tones at 5,000 Hz (5,000 Hz treatment); and (4) those exposed to silent treatment (no sounds playback for control). For 200 Hz treatment, we applied sounds with three sound pressure levels: 50 dB, 65 dB, and 100 dB, depending on experimental aims, whereas we only used that at 100 dB for the bee and 5,000 Hz treatments.

All types of sounds were collected from online source (bee sound; https://www.youtube.com/watch?v=AgY_3xLU4J0; 200 Hz sound; https://www.youtube.com/watch?v=xvF0iMiIOUU; 5,000 Hz sound; https://www.youtube.com/watch?v=cx1VQISKvhc): an Apple iPhone SE and a speaker (Peerless by Tymphany, XT25SC90-04) were used to play back all sounds. All the following experiments were conducted in 2025.

#### Nectar sampling and measurement

Within 10 minutes after each treatment, approximately 1.00 µL of nectar was collected from experimental flowers using micro-capillary tubes and diluted threefold for *R*. × *pulchrum* nectar. For *L. amplexicaule*, we collected 0.50 µL of nectar, which was measured directly without dilution. Sugar concentration was measured using calibrated Bellingham-Stanley low-volume Eclipse refractometers (0–50 Brix), which are accurate for volumes as low as 0.2 µL. The sugar concentration for each *R*. × *pulchrum* sample was calculated as the measured value multiplied by three.

#### Acoustic stimuli experiment

Firstly, for a single trial of each treatment, we selected a group of 4 or 6 flowers from a single individual of *R*. × *pulchrum*. In each trial, sound playback was presented to a group of experimental flowers by moving a speaker from flower to flower for 3 minutes, positioning it 15 cm from each flower for ca. 10 seconds before moving to the next, then returning to the first flower to complete the cycle. This protocol mimicked a pollinator hovering around the flower group (Veits *et al*. 2019). As a result, each experimental flower was exposed to sound for approximately 33.8 ± 0.3 seconds on average. We used 200 Hz treatment at 65 dB due to the output limitations of our audio system at this stage of the experiment. The experiments were conducted outdoors at around 20°C under natural environmental conditions. Fully opened, healthy flowers were selected for the experiments. For *R*. × *pulchrum*, flowers from the same plant were used in different treatments on the same day. Nectar was collected from two flowers and pooled for a single measurement sample to compensate for insufficient nectar volume per flower in *R*. × *pulchrum*. The experiment was conducted for three days (on 27 April and 5 May in the Aotani-cho park (N34.71, E135.21) and on 11 May in a Kobe University campus (N34.73, E135.23). We tested 149 samples from 298 flowers from 15 plants in total (36 samples from 72 flowers for silent treatment, 38 samples from 76 flowers for bee treatment, 38 samples from 76 flowers for 200 Hz treatment, 37 samples from 74 treatment for 5000 Hz).

In *L. amplexicaule*, we compared sugar concentrations only between bee and silent treatments. For a single trial of each treatment, we exposed all flowers from 2–3 individuals to sound playback of bee or silent control by moving a speaker from flower to flower for 3 minutes, following the same protocol as in the *R*. × *pulchrum* experiment. *Lamium amplexicaule*, nectar volume from each individual flower was always insufficient for a single measurement, so we pooled nectar from several flowers until 0.50 µL was collected for each sample. We conducted the experiment for two days (19 and 26 April) in the Kobe University campus. We examined 25 samples from 456 flowers (10 samples from 227 flowers for silent treatment, and 15 samples from 238 flowers for bee treatment) in this experiment.

#### Acoustic pressure experiment

To examine the effects of sound pressure level on 200 Hz treatment in *R. × pulchrum*, we subsequently conducted two field experiments. Flowers were exposed to artificial sinusoidal tones at 200 Hz with two sound pressure levels (50 dB and 65 dB), and compared sugar concentrations of these flowers with those exposed to silent treatment to examine an acoustic sensitivity threshold in *R. × pulchrum*. Additionally, we compared sugar concentrations between experimental flowers exposed to 200 Hz treatment at 100 dB and silent treatment as a follow-up experiment. The sound treatments were applied in the same manner as in the acoustic stimuli experiment. We conducted the experiments for two days: 50 dB and 65 dB treatments on 27 April in the Aotani-cho park, and 100 dB treatments on 11 May in the Kobe University campus. In total, 200 flowers were examined (40 samples from 80 flowers for silent treatment, 80 samples from 40 flowers for 50 dB treatment, 80 samples from 40 flowers for 65 dB treatment, and 80 samples from 40 for 100 dB treatment).

#### Floral organ removal experiment

All following organ-removed flowers in this experiment were exposed to an artificial sinusoidal tone at 200 Hz at 65 dB or silent treatment. We prepared four types of flowers without specific floral organs, flowers with petal removed (petal-removed); those with stamens and pistils removed (stamen-pistil-removed); those with stamens removed (stamen-removed); those whose pistils were removed (pistil-removed). We examined 40 flowers for each treatment for two days (27 April and 5 May): totally 160 samples from 320 flowers were treated (four types of flowers *×* two sound treatment *×* 20 samples from 40 flowers) in the Aotani-cho park.

### Statistical analysis

We analyzed the effects of sound treatments using generalized linear mixed models (GLMMs) with Gaussian error distribution and an identity-link function. For the acoustic stimuli experiment analysis, the response variable was nectar sugar concentration of each sample and the explanatory variable was sound treatments (silent/bee/200 Hz (65 dB)/5,000Hz for *R. × pulchrum* and silent/bee for *L. amplexicaule*). To account for potential unexamined variation across experimental days, observation date identity was included as a random effect.

For the acoustic pressure experiment GLMM analyses, the response variable was nectar sugar concentration of each flower, and the explanatory variable was sound treatments (silent/200 Hz at 100 dB or silent/200Hz at 50 dB/200Hz at 50 dB). To account for potential variation across experimental days, observation date identity was included as a random effect.

Finally, for the floral organ removal analysis, the explanatory variables were sound treatments (silent/200 Hz), removal treatments (petal/pistil-stamen/pistil/stamen), and their interaction whereas the response variable was nectar sugar concentration of each experimental flower. We included observation date identity as a random effect in the GLMM.

We tested for significant differences in each treatment effect relative to the silent control using a Wald test. We also conducted post hoc tests using Tukey’s method with a significance level set at 5%. All analyses were performed using the glmmTMB package (Fournier *et al*. 2012) and multiple comparisons using the multcomp package in R software (ver. 4.5.0; R Core Team, 2018).

## Results

### Acoustic stimuli and pressure experiments

Bee and 200 Hz (65dB) treatments significantly increased sugar concentration by ca. 8.7% compared to silent and 5,000 Hz treated in *Rhododendron* flowers (Fig. 1a). In contrast, no apparent increase in sugar concentration was found in the 5,000 Hz treatment compared to the silent treatment (Fig. 1a).

**Figure 1.**
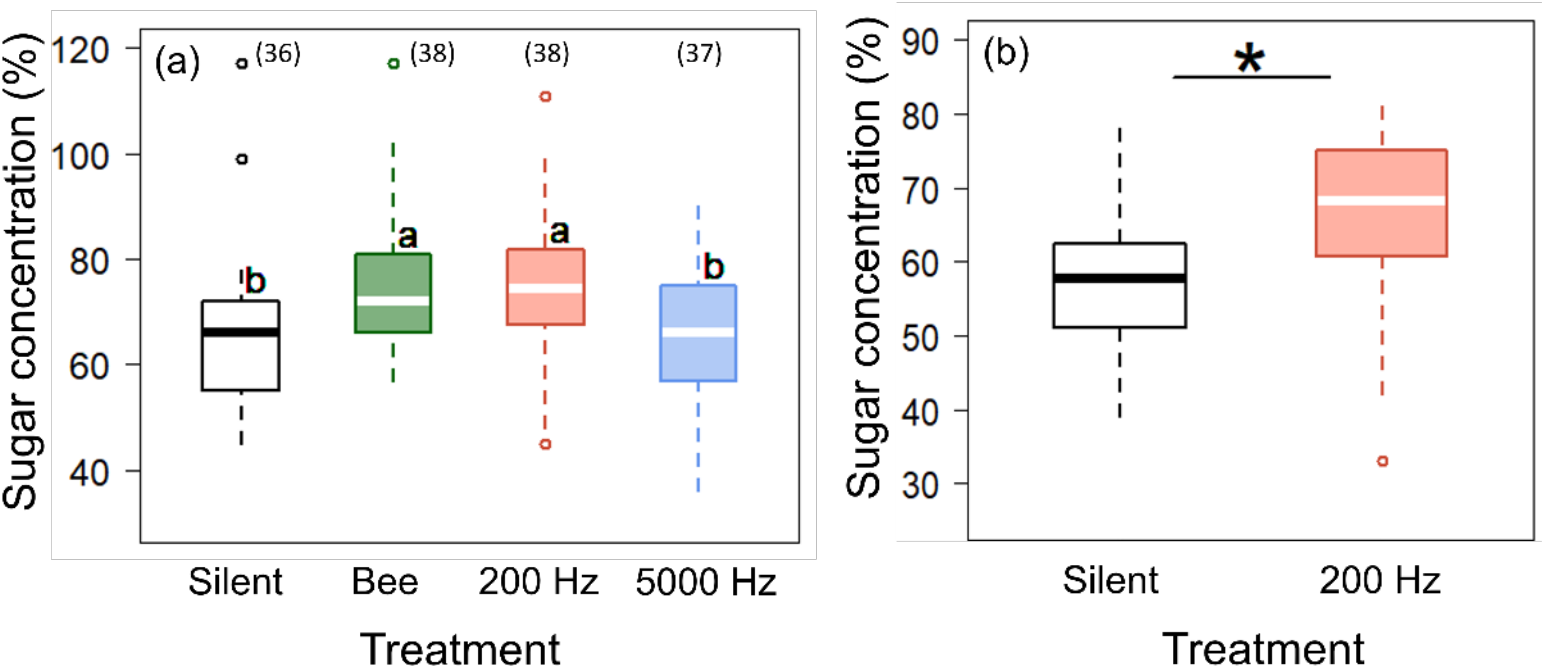
Comparisons of nectar sugar concentrations in *Rhododendron* × *pulchrum* flowers among sound exposure treatments: (a) silent (white), bee sound (100 dB, green) and artificial sound at 200 Hz (65 dB, red) and at 5000 Hz (100 dB, blue) (sample size); (b) silent and 200 Hz (100 dB). The central bars indicate the medians in the boxplots, and different alphabets in and an asterisk show significant differences (*p* < 0.05).

Furthermore, *Rhododendron* flowers exposed to 200 Hz (100 dB) sounds exhibited ca. 9.1% higher nectar sugar concentration than silently treated flowers in the follow-up acoustic pressure experiment (Fig. 1b). Additionally, we found no effects of 50 dB sounds on nectar sugar concentration increase, whereas 65 dB sounds significantly promoted higher nectar sugar concentration compared to silent treatment, indicating a sound response threshold between 50 and 65 dB (Fig. 2).

**Figure 2.**
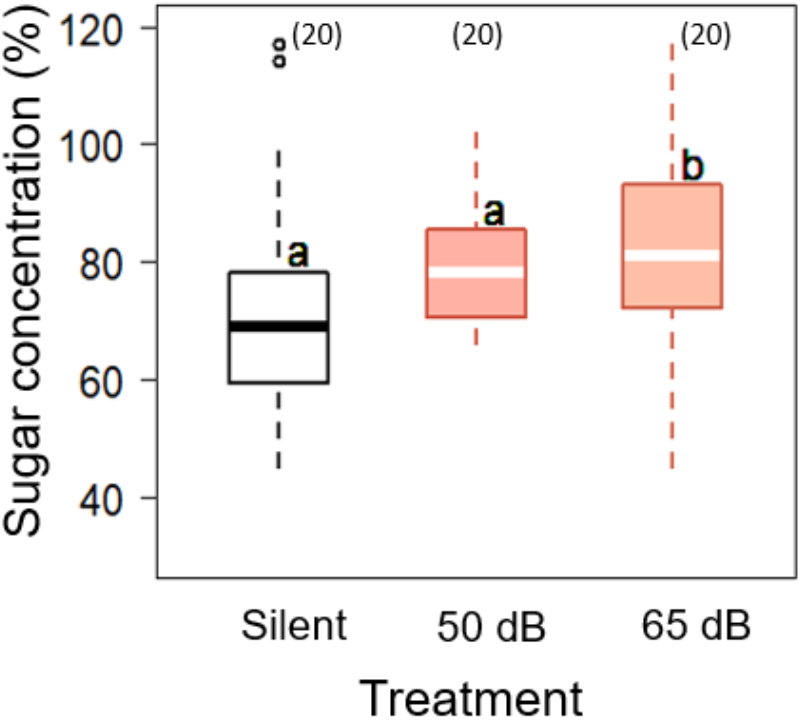
Comparison of nectar sugar concentrations in *R*. x *pulchrum* flowers among silent (white) and 200 Hz sound (red) treatment with different sound pressure (50 dB and 65 dB) (sample size). The central bars indicate the medians in the boxplots, and different alphabets show significant differences (*p* < 0.05).

In *L. amplexicaule* flowers, bee treatment significantly increased nectar sugar concentration (ca. 6.5% increase) compared to silent treatment as observed in *Rhododendron* flowers (Fig. 3).

**Figure 3.**
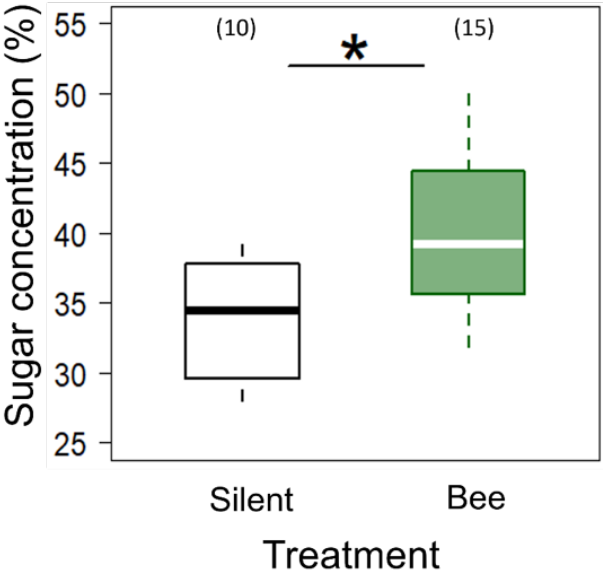
Comparison of nectar sugar concentrations in *Lamium amplexicaule* flowers between silent (white) and bee sound (100 dB, green) treatment (sample size). The central bars indicate the medians in the boxplots, and an asterisk show significant differences (*p* < 0.05).

### Floral organ removal experiment

We found no sugar concentration increases in petal-removed, stamen-pistil-removed, and stamen-removed flowers exposed to 200 Hz treatment compared to silent treatment, whereas pistil-removed flowers exhibited significantly increased sugar concentration when exposed to 200 Hz treatment compared to when silent treatment (Fig. 4).

**Figure 4.**
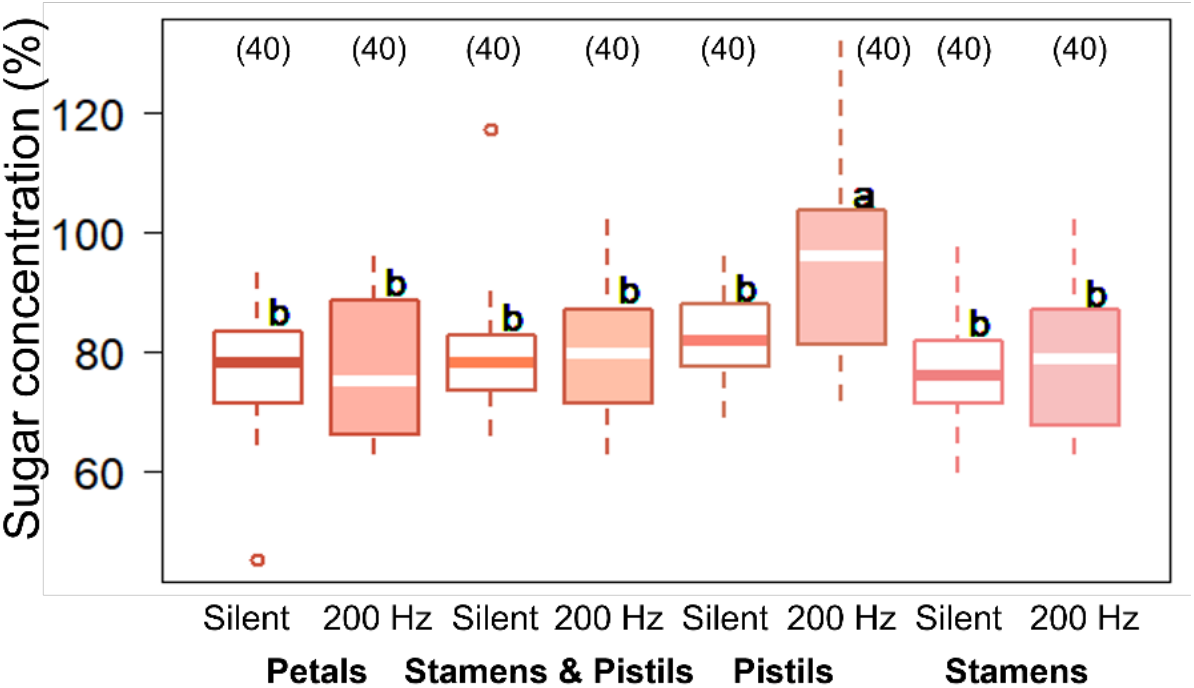
Comparison of nectar sugar concentrations in *R*. x *pulchrum* flowers between silent (white) and 200 Hz sound (65 dB, red) across different flower organ removal treatment: petal removal, stamens and pistils removal, pistils removal only and stamens removal only (sample size). The central bars indicate the medians in the boxplots, and different alphabets show significant differences (*p* < 0.05).

## Discussion

Our results demonstrate that flowers can perceive airborne bee sounds and adjust nectar sugar concentration accordingly in melittophilous *R. × pulchrum*. We also confirmed that exposure to pollinator-like frequencies (a synthetic 200 Hz tone at ≥ 65 dB) significantly increased nectar sugar concentration, whereas the silent control and high-frequency (5,000 Hz) treatments had no effect on sugar concentration increase in *R. × pulchrum*. Furthermore, the sensitivity of this response appeared to depend on sound pressure level, as we found a sound perception threshold between 50 and 65 dB, suggesting that the distance between pollinators and flowers is a critical factor in acoustic communication. The removal of petals and stamens eliminated this nectar response, suggesting that these floral structures are essential for acoustic perception. These findings support the idea that plants can sense pollinator sounds and rapidly modulate floral rewards to attract them. A similar response to bee sounds was also observed in *L. amplexicaule*, indicating that this ability may be widespread among bee-pollinated plant taxa, as an adaptation for bee attraction.

Rapid enhancement of nectar sugar concentration in response to pollinator sounds may be adaptive in maximizing pollinator revisitation. By providing high-quality rewards, flowers should be favored by large social bees such as bumble bees and honey bees, which discriminate differences in sugar concentrations among flowers and preferentially revisit flowers offering higher rewards (Greggers & Menzel, 1993; Chittka *et al*. 1999; Nardone *et al*. 2013; Pamminger *et al*. 2019). This nectar enrichment strategy enables plants to minimize investment when pollinators are absent while enhancing pollination success when reliable pollinators are present. However, bees’ foraging decisions may be influenced by trade-offs between nectar concentration, volume, and intake efficiency, rather than concentration alone (Nardone *et al*. 2013). Because we measured only sugar concentration due to technical limitations in nectar collection, further studies are warranted to examine this issue.

Moreover, whether this phenomenon extends beyond melittophilous plants remains uncertain. Lepidopterans, such as butterflies, also show a general preference for flowers with higher sugar concentration, suggesting that a similar sound response may be found in lepidopteran-pollinated flowers. The fact that both *R*. × *pulchrum* and *L. amplexicaule* examined in this study, as well as *O. drummondii* are sometimes visited by butterflies and hawkmoths, raises the possibility that sound-induced nectar modulation may be found in flowers visited by other pollinator groups, although further empirical studies will be necessary to test this hypothesis.

In *O. drummondii*, the petals have been suggested to function as sound-sensing organs in flowers (Veits *et al*. 2019). In our study, both the stamens and petals play an essential role in detecting bee sounds in *R*. × *pulchrum*. These organs may discriminate relevant acoustic cues from background noise in natural soundscapes where wind, birds, and other insects generate a wide range of frequencies, suggesting a high degree of signal specificity. Petals usually exhibit multifunctionality, such as attracting pollinators, protecting reproductive organs, and functioning as landing platform for pollinators (Endress *et al*. 2006; Katsuhara *et al*. 2017), while stamens are primarily responsible for pollen production and transfer. Our findings show that these floral organs have an additional function in perceiving flying pollinator sounds. However, our data are insufficient to discuss the detailed mechanisms by which these organs sense bee and bee-like sounds; therefore, further biomechanical studies are required to elucidate this process.

Despite the limitations noted above, we confirmed the generality of the findings by Veits *et al*. (2019) using two melittophilous species from different lineages than *Oenothera*. We suggest that further experimental studies are required to expand this generality in plants with other pollinator syndromes, and to elucidate the biomechanical, cellular, and molecular bases of sound detection in flowers. These processes potentially involve mechanoreceptors, calcium signaling, and rapid gene expression changes, which remain key challenges.

## Author contribution

Conceptualization, KS; methodology, AT, AU, GSH,KS; validation, AT, AU, GSH,KS; formal analysis, AU, GSH, KS; investigation, AT, KS; resources, AT; data curation, AU, GSH, KS; writing—original draft preparation, KS; writing—review and editing, AT, AU, GSH,KS; visualization, AU, GSH, KS; supervision, AT, AU; project administration, AT; funding acquisition, AT, KS. All authors have read and agreed to the published version of the manuscript.

## Acknowledgement

The authors are also grateful to Prof. Nobuko Ohmido for providing the experimental site. The authors thank the JST Global Science Campus and JST Science and Technology Challenge Program for Next Generation (ROOT Program) for supporting this study.

## Notes

### Competing Interest Statement

The authors have declared no competing interest.

